# Legacy copper/nickel mine tailings potentially harbor novel iron/sulfur cycling microorganisms within highly variable communities

**DOI:** 10.1101/2024.01.24.577127

**Authors:** Molly Chen, Daniel S. Grégoire, Jeffrey G. Bain, David W. Blowes, Laura A. Hug

## Abstract

The oxidation of sulfide-bearing mine tailings catalysed by acidophilic iron and sulfur-oxidizing bacteria releases toxic metals and other contaminants into soil and groundwater as acid mine drainage. Understanding the environmental variables that control the community structure and metabolic activity of endogenous microbes living in tailings (especially the abiotic stressors of low pH and high dissolved metal content) is crucial to developing sustainable bioremediation strategies. We determined the microbial community composition along two continuous vertical gradients of Cu/Ni mine tailings at each of two tailings impoundments near Sudbury, Ontario. 16S rRNA amplicon data showed high variability in community diversity and composition between locations, as well as at different depths within each location. A temporal comparison for one tailings location showed low fluctuation in microbial communities across 2 years. Differences in community composition correlated most strongly with pore-water pH, Eh, alkalinity, salinity, and the concentration of several dissolved metals (including iron, but not copper or nickel). The relative abundances of individual genera differed in their degrees of correlation with geochemical factors. Several abundant lineages present at these locations have not previously been associated with mine tailings environments, including novel species predicted to be involved in iron and sulfur cycling.

## IMPORTANCE

Mine tailings represent a significant threat to North American freshwater, with legacy tailings areas generating acid mine drainage (AMD) that contaminates rivers, lakes, and aquifers. Microbial activity accelerates AMD formation through oxidative metabolic processes but may also ameliorate acidic tailings by promoting secondary mineral precipitation and immobilizing dissolved metals. Tailings exhibit high geochemical variation within and between mine sites, and may harbor many novel extremophiles adapted to high concentrations of toxic metals. Characterizing the unique microbiomes associated with tailing environments is key to identifying consortia that may be used as the foundation for innovative mine-waste bioremediation strategies. We provide an in-depth analysis of microbial diversity at four copper/nickel mine tailings impoundments, describe how communities (and individual lineages) differ based on geochemical gradients, predict organisms involved in AMD transformations, and identify taxonomically novel groups present that have not previously been observed in mine tailings.

## BACKGROUND

Mine tailings are the fine-grained residues left as waste from selective flotation of sulfide minerals, as part of the extraction process of valuable metals from ores. Sulfide-rich tailings stored in subaerial impoundments are a major source of groundwater contamination due to release of acid mine drainage (AMD). When sulfidic minerals (*e.g.*, pyrite, FeS_2_) are exposed to oxygen and water, the oxidation of ferrous iron (Fe^II^) and sulfur generates high amounts of Fe^II^ SO_4_^2-^ and H^+^. The Fe^II^ release by sulfide mineral oxidation may be further oxidized to Fe^III^, which contributes to the oxidation and dissolution of additional pyrite and other metal-bearing minerals. The overall effect of these reactions includes the formation of acidic porewater with high concentrations of heavy metals. Release of AMD to natural environments and freshwater is detrimental to natural ecosystems and human health (reviewed in Simate & Ndlovu, 2014).

Microbial activities can accelerate the rate of AMD formation by hundreds to millions of times compared to abiotic oxidation alone (Nordstrom et al., 2015; Singer & Stumm, 1970). Acidophilic, chemolithotrophic bacteria such as *Acidithiobacillus ferrooxidans* (Leduc & Ferroni, 1994; Wang et al., 2019) obtain energy through the oxidation of iron and sulfur compounds, releasing additional H^+^, SO_4_^2-^, and regenerating the Fe^III^ ion. While these organisms are detrimental in the context of tailings waste management, they have also been extensively studied for industrial bioleaching applications to extract valuable metals from tailings (Falagán et al., 2017; Valdés et al., 2008).

Microbial consortia isolated from AMD environments also include acidophilic sulfate reducers that precipitate metal ions from solution as metal sulfides (Johnson et al., 2013), and species that immobilize metal ions through bioaccumulation or biosorption (Anusha et al., 2021; Bautista-Hernández et al., 2012; Mohamad et al., 2012; Tsuruta, 2011); all of these processes serve as potential remediation approaches to remove metals from tailings porewater and mitigate the negative environmental and human health impacts of mine wastes. Physical remediation strategies may also be employed to reduce acidity and metal leaching of tailings, such as by the application of cover layers (*e.g.*, fine-grained silt and clay or organic carbon) that limit oxygen and water ingress into the tailings and facilitate the growth of vegetation, (Aubertin et al., 2016; Pakostova et al., 2022a; Park et al., 2019; Peppas et al., 2000). Vegetation (phytoremediation) and organic matter in organic covers also increase microbial community diversity of the tailings by promoting the establishment of organoheterotrophs and nitrogen-fixing rhizobacteria (increasing plant growth, nutrient turnover, and ecosystem productivity) (Asemaninejad et al., 2020; Romero et al., 2021).

Many studies have described microbial genera common across varied AMD environments, such as the acidophilic iron and/or sulfur oxidizing *Acidithiobacillus*, *Leptospirillum*, and *Sulfobacillus* spp. (reviewed in Baker & Banfield, 2003; Méndez-García et al., 2015). However, mine tailings can be diverse in composition depending on the metals targeted for extraction, resulting in substantial differences in the extent of oxidation and acidification and concentrations of contaminants. As a result, the microbial community compositions found within mine tailings are also highly heterogenous. pH is an important driving factor in the relative abundance of several lineages, with Proteobacteria generally dominant in slightly to moderately acidic tailings (pH 4-7) and Euryarchaeota (especially *Ferroplasma* spp.) more abundant in highly acidic tailings (pH <3) (Chen et al., 2013; Jones et al., 2017; Liu et al., 2014). The concentrations of metal ions (including Fe, Cu, Ni, and trace elements such as Cd, Hg, and U) select for organisms with resistance mechanisms against these contaminants (Bondici et al., 2013; Jones et al., 2017; Liu et al., 2014; Yan et al., 2020). Mine tailings also exhibit vertical stratification, with distinct zones of oxidation, neutralization, and unaltered tailings, heterogeneous physical characteristics (*e.g.,* particle size) and temporal variability in moisture content; all of which influence the microbial guilds dominating each microenvironment (Diaby et al., 2007).

An accurate view of the specific microbiomes associated with different tailings environments is key to developing sustainable and cost-effective measures to remediate mine wastes (*e.g.*, bioreactors with engineered sulfate-reducing consortia, *in-situ* remediation via iron-oxidizing consortia in aerobic wetlands) (Johnson & Hallberg 2005). Tailings are highly selective environments that may be host to unexplored phylogenetic radiations of bacteria and archaea that have evolved unique adaptations to survive in heavy metal contaminated areas. Therefore, tailings provide an excellent starting point to identify AMD-tolerant microbes that can be used to improve the efficiency of engineered bioreactors and *in-situ* remediation strategies. Recent advances in high-throughput sequencing methods such as 16S rRNA amplicon sequencing have enabled the high-resolution examination of the taxonomic diversity of AMD microbiomes (reviewed in Lukhele et al., 2019).

In this study, we compared microbial community diversity across vertical and horizontal transects of tailings associated with two Ni/Cu mining sites in the Sudbury Basin region of Ontario, Canada. 16S rRNA amplicon sequencing was used to characterize community composition across 3-5 m depth transects at four tailings locations. A temporal comparison was made for one tailings location to assess community stability over time (2 years). Our objectives were to identify the main geochemical factors influencing microbial diversity and variability between locations, predict the contribution of microbial guilds to iron/sulfur cycling within tailings, as well as to explore microbial novelty within the context of mine tailings environments.

## RESULTS AND DISCUSSION

### Sampling site and tailings core characteristics

The Sudbury Basin is a major geological structure in the Canadian Shield. Formed by a meteorite impact (French, 1967), it consists of mainly Fe-Ni-Cu sulfide deposits (*e.g.,* pyrrhotite, chalcopyrite and pentlandite), which may be enriched in precious metals (Molnár et al., 1999).

Tailings stored at two mines owned and operated by Glencore Sudbury Integrated Nickel Operations (Sudbury INO) were sampled for characterization of microbial community diversity and function: the Strathcona Waste Water Treatment System (SWWTS), which received tailings from 1968 – 2012 (Bain et al., 1998; Bain & Blowes, 2013) and Nickel Rim North tailings area, which received tailings from 1953 – 1958 (Johnson et al., 2000). Operations at Sudbury INO are focused primarily on extracting and processing nickel and copper, with cobalt and precious metals as by-products (Glencore Canada, 2022).

At Strathcona, two locations, Moose Lake 25 (ML25) and ML34, were chosen to assess spatial (vertical) variation. The sulfide-rich (9-18 wt. % S; Blowes et al., 1999), coarse-grained tailings at ML25, which have been exposed to atmospheric weathering since deposition decades ago, are extensively oxidized. The porewater at ML25 is low pH (pH < 5) and contains high concentrations of contaminants (McAlary, 2020). In contrast, the high sulfide tailings at ML34 are overlain by a 2 m layer of desulfurized tailings (DST) cover of < 1.4 wt. % S mainly as pyrrhotite, and a 50 cm organic carbon cover composed of a mixture of a mixed municipal compost amended with biosolid fertilizer. The cover materials are not potentially acid generating, and the fine-grained DST cover maintains a high moisture content, limiting O_2(g)_ diffusion. The pH is circumneutral and concentrations of dissolved metals are lower than at ML25 (McAlary, 2020; Pakostova et al., 2022).

At Nickel Rim North (NR), locations NR18 and NR3 were selected for comparison. The unoxidized tailings contain on the order of 3 wt. % S mainly as pyrrhotite (Johnson, 1993). Contaminant profiles are similar at these two locations, although NR3 has shallow water-table elevation and the shallow tailings have a high moisture content, leading to a shallower depth of oxidation and lower concentrations of dissolved metals than at NR18 (Johnson, 1993). NR18 also has a shallow (< 10 cm) organic soil, deposited in the early 1990s (Johnson, 1993).

In 2021, tailings core samples extracted from each location were collected in lengths of aluminum pipe (with inner diameters measuring 5.08 and 7.62 cm) driven into the tailings using the technique of Starr and Ingleton (1992). The core samples were subsectioned in lengths of 10 cm (except for the ends of each core, which were 10-17 cm), generating a total of 112 samples (Supplemental Table 1). A core collected, frozen and processed in the same way from NR18 in 2019 was used for comparison to 2021 samples (Supplemental Table 1).

### 16S rRNA sequencing statistics

Out of a total of 7,610,282 raw reads (ranging from 161 to 217,042 per sample), 6,100,701 were kept after filtering, denoising, merging, and detection of chimeric reads. From these, 11,116 amplicon sequence variants (ASVs) were identified. The median ASV frequency per sample was 50,912, though the range varied from 141-199,300, reflecting the variability in community composition within samples. The median frequency of each ASV across all samples was 29, with a maximum frequency of 546,045. ASVs with a total frequency of 100 or less were generally confined to 1 or 2 samples, while higher abundance ASVs were present across multiple cores and/or multiple depth ranges.

### Alpha diversity

Plots comparing of Faith’s phylogenetic distances, Shannon indices, and Observed ASVs of samples (grouped by location) showed consistent trends across the four cores (Figure 1). Samples from ML25 consistently scored low on α-diversity metrics across all depths, although the specific taxonomic composition of samples was highly variable when comparing upper and lower depths (discussed in the Community Composition section). The α-diversity metrics of NR3 samples were overall similar to ML25, although the deeper fractions of the core tend to have higher diversity (Figure 1). It is possible that the deposition of NR3 tailings on top of native vegetation in the 1950s, the presence of nearby plants, and periodic flooding events (Johnson, 1993) account for the higher diversity at greater tailings depths, supported by the increased DOC concentrations in these samples (Table 1).

**Figure 1:**
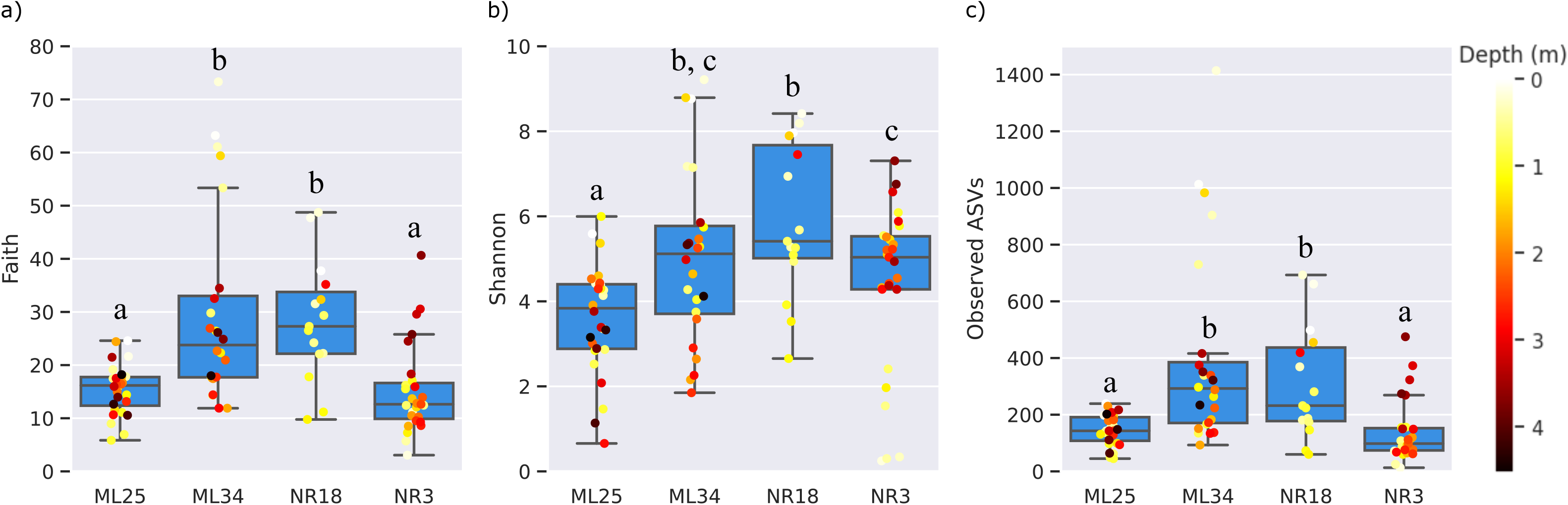
*α*-diversity metrics for all sampling locations and depths. Faith’s phylogenetic distances (a), Shannon diversity indices (b), and observed ASVs (c) were calculated with QIIME2 using a rarefied sampling depth of 12,000 reads. Individual samples overlaid on box plots are colored according to the vertical depth below ground level. Significant differences (Kruskal-Wallis, p<0.05) between locations are designated by different letters above each boxplot.

ML34 and NR18 showed significantly higher α-diversity metrics compared to ML25/NR3, which we attribute to the cover layers and vegetation present at these locations (Londry & Sherriff, 2005; Pepper et al., 2012), and is especially evident in the fact that ML34 diversity was highest in the samples closest to the surface. A similar trend was seen for NR18, but due to low read numbers in the deeper NR18 sections, most samples below 2 metres were excluded after rarefying to a minimum sampling depth of 12,000, hindering a full comparison.

### Beta diversity

Samples did not appear to discretely cluster by location in any of the β-diversity ordinations (Figure 2a, Supplemental Figures 1a & 2a), indicating that the variability between samples at the same location was greater than the overall differences between locations.

**Figure 2:**
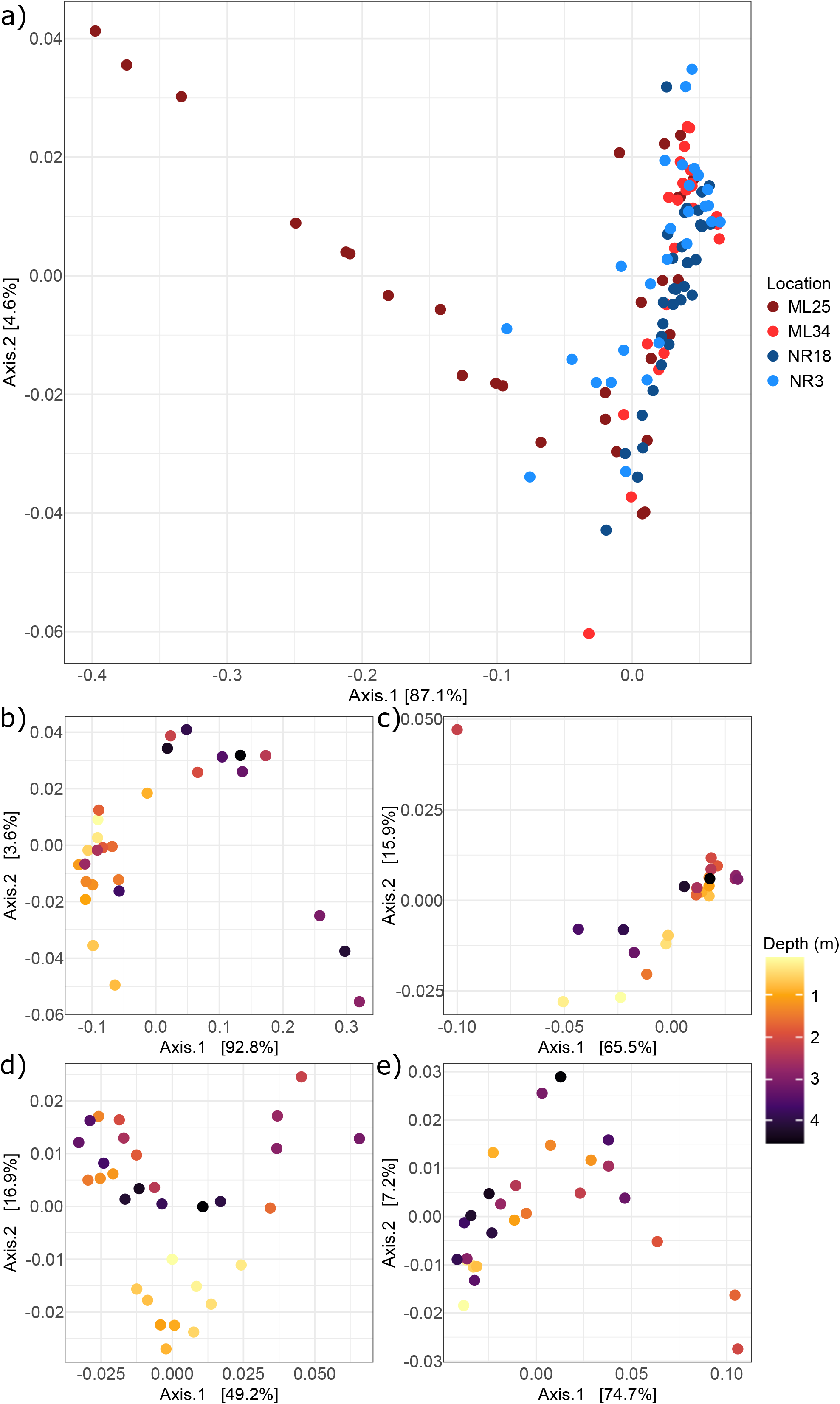
Weighted unifrac PCoA plot for. **a)** all sampling locations combined, **b)** ML25, **c)** ML34, **d)** NR18, and **e)** NR3. Samples in plots b-e are colored by depth.

The weighted unifrac principal coordinate analysis (Figure 2a) was the most effective in mapping distances between samples, with the two principal axes explaining 91.7% of variation in the data. ML25 samples showed particularly high separation along the primary axis, which may reflect the much broader range of contaminant concentrations along the core, such as dissolved Fe, Ni, Cu, and SO_4_^2-^ (Supplemental Data File 1). Analysis of individual locations showed that weighted unifrac PCoA was also best for mapping distances (Figure 2b-e), with the principal axes explaining 66.1 – 96.4% of the variation. There was some separation of samples along one or both axes based on depth, which was more apparent in the Bray-Curtis and unweighted unifrac PCoA plots (Supplemental Figures 1b-e & 2b-e), suggesting the presence of environmental gradients along the depth of the core that influence community composition.

NMDS analysis (Figure 3) also did not show clustering of samples by location, though there was some separation between ML and NR samples along the secondary axis. Out of all geochemical variables mapped to the samples (Supplemental Data File 1), the ones significantly correlating with community composition included pH, Eh, alkalinity, and salinity (inferred by electrical conductivity (EC), and dissolved Na, Cl, and K). It was expected that major community shifts occur across these environmental gradients, as microorganisms segregate into niches based on tolerance to selective forces (*e.g.*, stress from low pH or high salinity) (Sardinha et al., 2003; Zhalnina et al., 2015; Zhang et al., 2019) and metabolic potential (influenced by redox gradients and oxygen availability) (Chen et al., 2017; Lipson et al., 2015; Pett-Ridge & Firestone, 2005). Pore-gas oxygen measurements were taken at ML25, ML34, and NR18 tailings, but not included in the statistical analyses due to limited data points (Supplemental Figure 4). However, the iron-dominated redox potential (Eh) of tailings environments can be used as a proxy to infer the metabolic capacity of samples (Supplemental Data File 1); with aerobic respiration only supported at highly positive Eh values (above 300-500 mV) (Søndergaard, 2009), although this threshold increases with decreasing pH.

**Figure 3:**
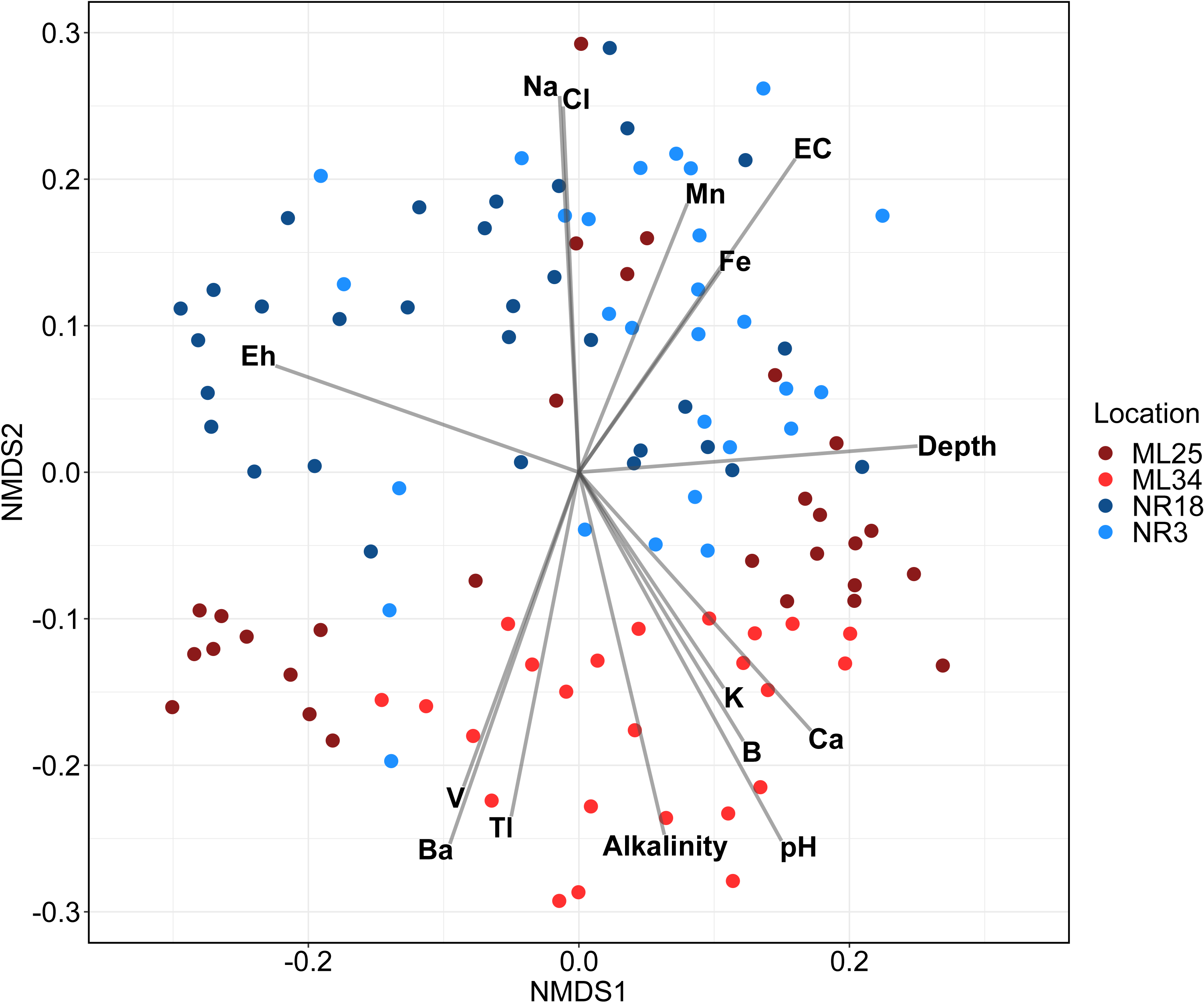
NMDS biplot of Bray-Curtis distances and environmental variables. Samples are colored by location. Environmental variables that significantly correlated with community dissimilarity (p < 0.05, Bonferroni corrected) were included. For a plot showing all variables, see Supplemental Figure 3.

Diversity also correlated with the concentrations of Fe, Mn, B, Ba, Ca, Tl, and V. Ferrous and ferric iron availability is known to correlate with the abundance of iron oxidizing and reducing bacteria, such as members of the Actinobacteria and Gammaproteobacteria (Liu et al., 2014). Many iron oxidizing/reducing organisms can also oxidize/reduce manganese (de Vrind-de Jong et al., 1990; Thamdrup, 2000). As for the other elements, correlation between microbial diversity and concentration is more likely due to the co-variance between ion concentration and another geochemical factor, such as pH, rather than direct interactions between the elements and microorganisms (Supplemental Figure 5). For example, calcium (*i.e.,* Ca in Figure 3) concentration was strongly correlated with alkalinity (measured in mg/L CaCO_3_) (R=0.79). However, it is possible that selection for resistance to Tl and V toxicity may influence community composition (She et al., 2022; Zhang et al., 2019), as these elements are highly toxic at trace concentrations. In particular, vanadium concentrations in several samples from ML25 and ML34 far exceed the recommended safe limit for drinking water (0.05 mg/L) (Environment and Climate Change Canada, 2016) so selection for vanadium resistance mechanisms such as the reduction of V^V^ to V^IV^ would be relevant in these communities.

Interestingly, β-diversity did not significantly correlate with Ni or Cu concentrations, despite these elements being the major porewater contaminants present in these tailings besides Fe (with upper concentration limits of 1,166 mg/L for Cu and 561.6 mg/L for Ni) (Supplemental Data File 1). A possible explanation could be that Ni/Cu tolerance is widely distributed across many microbial taxa commonly found in mine tailings, and thus the overall variance in community composition is determined by other selective forces. Ni and Cu are both micronutrients required for the function of metalloenzymes and most prokaryotes encode homeostatic mechanisms to regulate intracellular concentrations (reviewed in Newsome & Falagan, 2020). Nickel efflux pumps such as RcnA have been identified in organisms isolated from metal contaminated environments (*e.g.*, *Cupriavidus metallidurans* strain CH34), as well as in non-Ni-resistant *E. coli* and *H. pylori*, with homologs found in other proteobacteria, cyanobacteria, and archaea (Macomber & Hausinger 2011). Similarly, extremophiles used in industrial biomining such as *A. ferrooxidans* and *S. metallicus* share the same ATPases and transporters for copper export (CopA and CusA) with other bacteria, archaea, and eukaryotes (Orell et al., 2010, Andrei et al., 2020; Antsotegi-Uskola et al., 2020). The presence of these pathways in Nickel Rim/Strathcona Mill tailings could be confirmed with multi-omic sequencing.

### Community composition

Multiple changes in the dominant phyla occur along the depth gradients for each sampled core (Figure 4). In general, upper layers are mainly comprised of *Proteobacteria* and *Acidobacteriota*, which is consistent with most studies on mine tailing communities (Chen et al., 2013; Pepper et al., 2012; Yan et al., 2020). *Desulfobacterota* and *Firmicutes* are abundant in deeper tailings; members of these phyla include anaerobic sulfate-reducing bacteria commonly found in anoxic layers of tailings (Ayangbenro et al., 2018; Gao et al., 2022; Korehi et al., 2014; Winch et al., 2009). *Actinobacteriota* were highly abundant in ML25 tailings below 2 m in depth (Figure 4), but not at the other locations. While *Actinobacteriota* are commonly found in mine tailings (especially metal tolerant *Arthrobacter* spp.) (Gao et al., 2021; Lopez et al., 2017; Xie et al., 2022), these organisms are usually obligate aerobes associated with the rhizosphere of metal hyper-accumulating plants. Therefore, we would expect to see them in the surface layers of vegetated tailings (e.g., ML34) as opposed to the deeper, ML25 tailings. The *Actinobacteriota* ASVs in ML25 appear to belong to novel clades not previously found in mine tailings; their classification is discussed in the next section.

**Figure 4:**
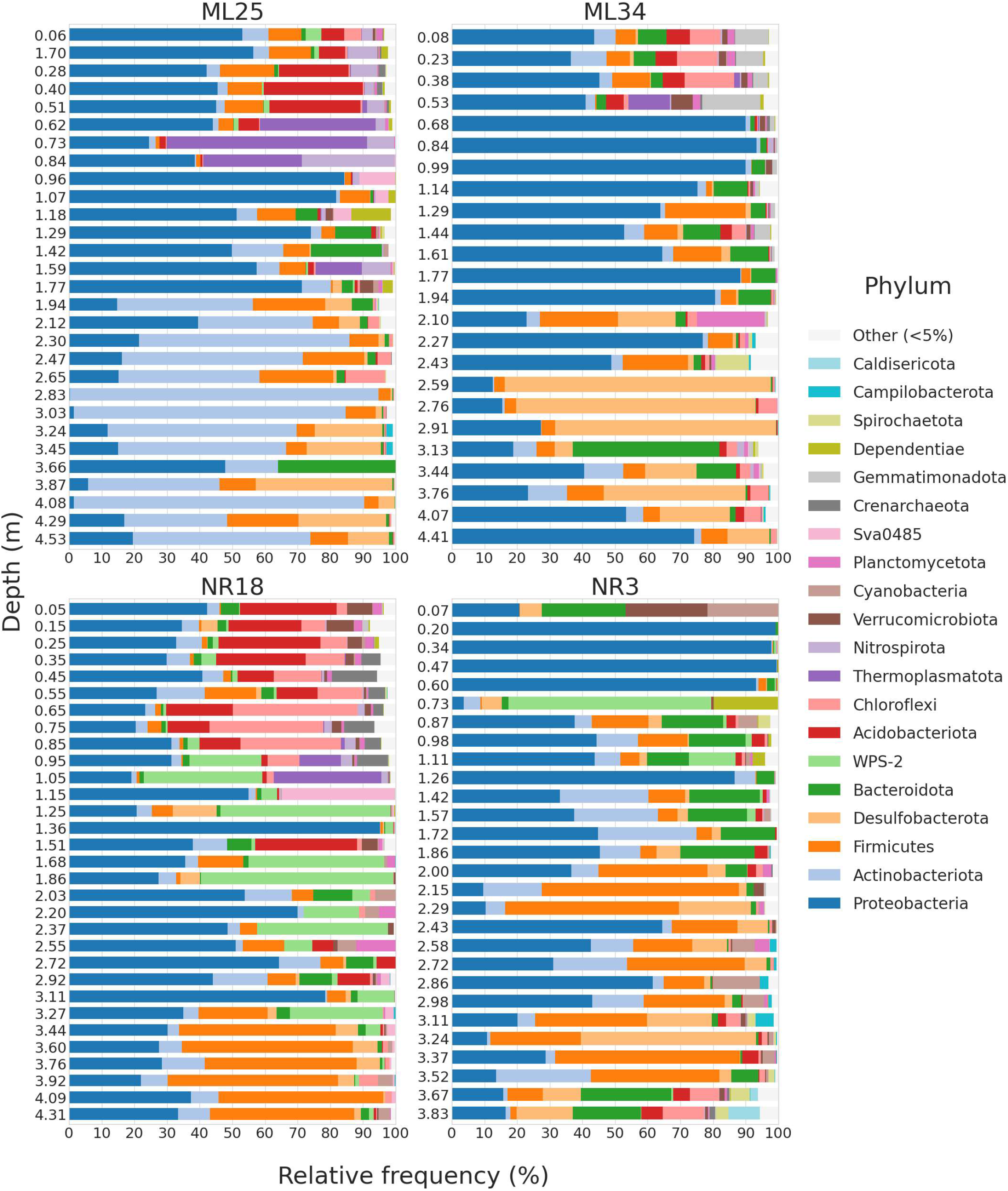
Phylum-level microbial community composition of mine tailings. Bars representing samples are grouped by location (ML25, ML34, NR18, NR3) and depth. Bars depict relative frequency of ASVs corresponding to all phyla present at a frequency of ≥5% in at least one sample. The bar widths (Y-axis) are not scaled based on the depth range covered.

The WPS-2 phylum is also novel in terms of its presence in mine tailings and seems to be unique to Nickel Rim samples (particularly at NR18 at depths between ∼1-3 m) (Figure 4). WPS-2 (or *Ca*. *Eremiobacterota*) is a recently described phylum, consisting of a wide range of metabolically diverse bacteria found in many types of terrestrial environments (Ji et al., 2021). Metagenomes isolated from Antarctic desert soil suggest some WPS-2 members are acidophilic (Ji et al., 2021). Another study found that the WPS-2 were abundant (with ASV relative frequencies of up to 24%) in the unvegetated soils of extinct iron-sulfur springs in British Columbia (Sheremet et al., 2020). Given that sampling occurred during near-freezing temperature periods and the acidic, iron/sulfur contaminated nature of AMD, it is not surprising that the WPS-2 could thrive in tailings environments. However, Ji et al. (2021) and Sheremet et al. (2020) found no evidence of microaerophilic or anaerobic metabolic capacity from genomic data in their studies, meaning the WPS-2 ASVs present at Nickel Rim likely represent organisms with novel metabolic capacity within this understudied lineage.

### Community change along geochemical gradients

A more detailed analysis of community composition changes (in terms of genus-level relative abundance) across tailings samples is shown in Figure 5a. The distribution of the most abundant genera (≥ 15% in at least one sample) was highly variable. Most populations did not show consistent abundance across samples and tended to be confined to and/or enriched in specific depth ranges.

**Figure 5:**
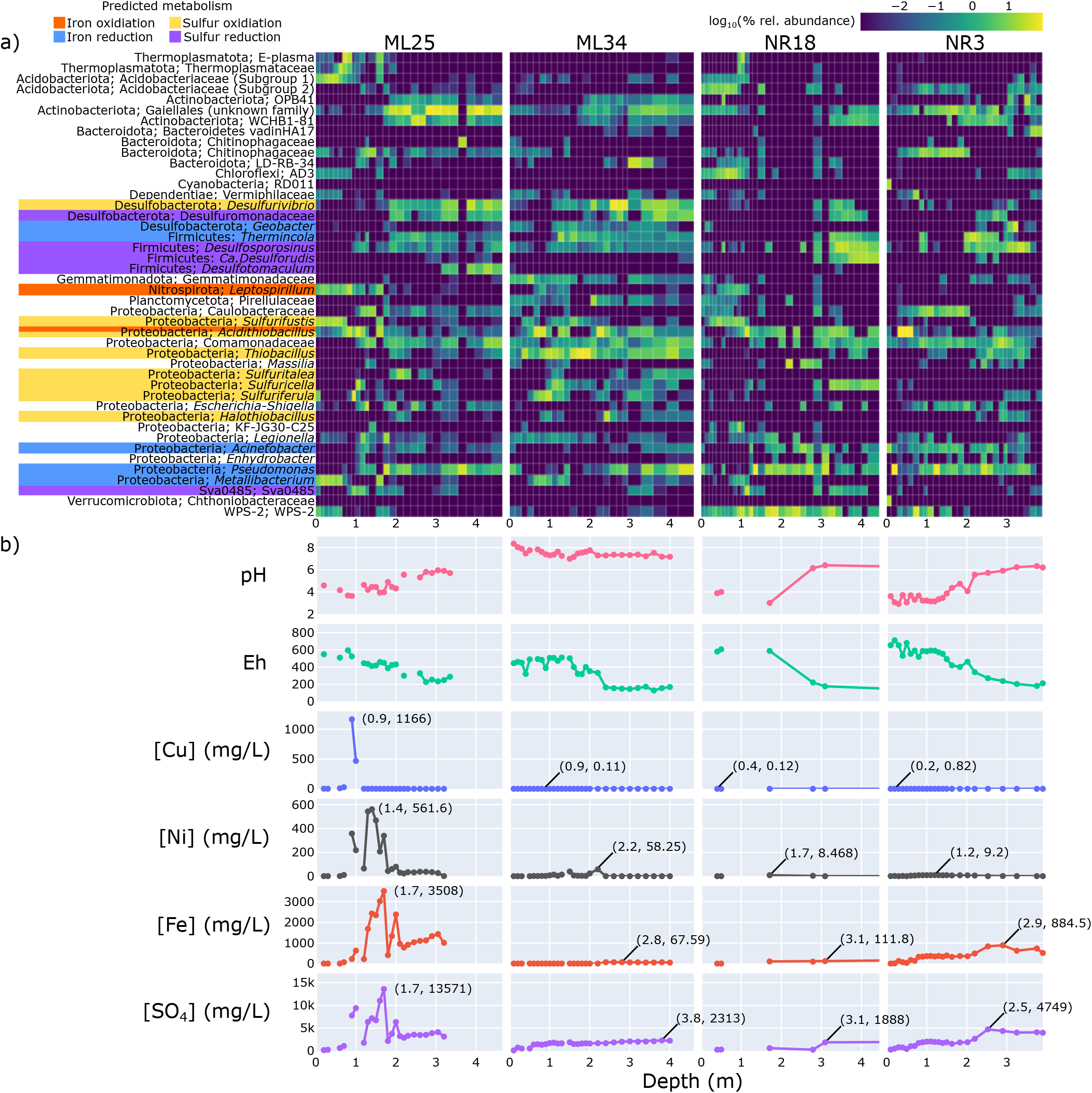
Distribution of bacterial and archaeal genera along tailings depth gradients and pore-water chemistry analyses. The heatmap (a) indicates the percent relative abundance of genera that were present at an abundance of at least 15% in at least one sample, across all depth intervals at each sampling location. The phylum and genus are labelled (if the genus was unknown, the family is shown instead). Relative abundances were log transformed, with 0% abundance values set to -2.999. Genera with predicted iron/sulfur cycling capabilities are indicated in color. The corresponding values for pH, Eh, and contaminant concentrations (Cu, Ni, Fe, SO_4_^2-^) are plotted in (b). For contaminant plots, the coordinates (depth in metres, concentration in mg/L) of the peak concentration value are indicated directly on the plot.

To track how variation in geochemistry along cores impacted the distribution of the lineages highlighted in Figure 5a, pH/Eh and the concentration of the most environmentally relevant contaminants (Fe, Ni, Cu, SO_4_) were linked to genera abundances (Figure 5b). The pH of ML25, NR18, and NR3 ranged from moderately to slightly acidic (∼3-6) wherein pH increased with depth, while the pH of ML34 was slightly above 7 and more stable across all depths. The general trend of peak Fe, Ni, and Cu concentrations was similar between locations: the highest copper concentrations were close to the surface, followed by nickel, and finally iron. This observation is consistent with previous hydrogeochemical assessments of Nickel Rim (Johnson et al., 2000). These differences are due to differences in the susceptibility of pyrrhotite (Fe_(1-x)_S), pentlandite ((Fe,Ni)_9_S_8_), and chalcopyrite (CuFeS_2_) to oxidation, and to the formation of secondary Fe oxyhydroxide phases (*e.g.,* goethite; αFeOOH) and the secondary Cu sulfide mineral covellite (CuS) (Johnson et al., 2000). Optical examination of mineral thin sections from Nickel Rim and similar tailings impoundments have shown that pyrrhotite is the most susceptible (least resistant) to oxidation, followed by pentlandite, chalcopyrite and pyrite (Lindsay et al., 2015). Other factors influencing the localization of dissolved metals include differences in metal sorption capabilities to soil/clay particles (competitive sorption of Cu ions is greater than Ni ions, reducing transport to lower depths) (Sheikhhosseini et al., 2013; Harter, 1992) as well as pH-dependent precipitation of Fe (oxy)hydroxides (which can sorb Ni) and covellite (CuS) (Johnson, 1993). Sulfate concentrations generally followed Fe, as both are released during the dissolution of sulfide minerals (see Supplemental Figure 5 for a summary of correlations between geochemical variables).

Correlations between abundance data (Figure 5a) and geochemistry (variables from Figure 5, plus Ni, Cu, and SO_4_) identified several significant connections (Figure 6). We included Cu and Ni in our analyses despite their concentrations not having a significant impact on overall community diversity because they are the main contaminants released from these tailings, and increased tolerance to these metals would be an asset for bioremediation-relevant organisms. Dissolved Cu concentrations were moderately correlated with the abundance of almost half of the dominant genera (21 of 44 genera, 48%). Ni showed comparatively fewer significant correlations (Figure 6). As with the NMDS analysis, it was difficult to determine which specific factors were directly interacting with microorganisms, due to the interconnected nature of environmental variables (Supplemental Figure 5). Correlation matrices were also individually calculated for each sampling location (Supplemental Figure 6), which differed from the pooled dataset (most notably, ML25 had much stronger R values due to a wider range of contaminant concentrations, contributing to stronger selective forces).

**Figure 6:**
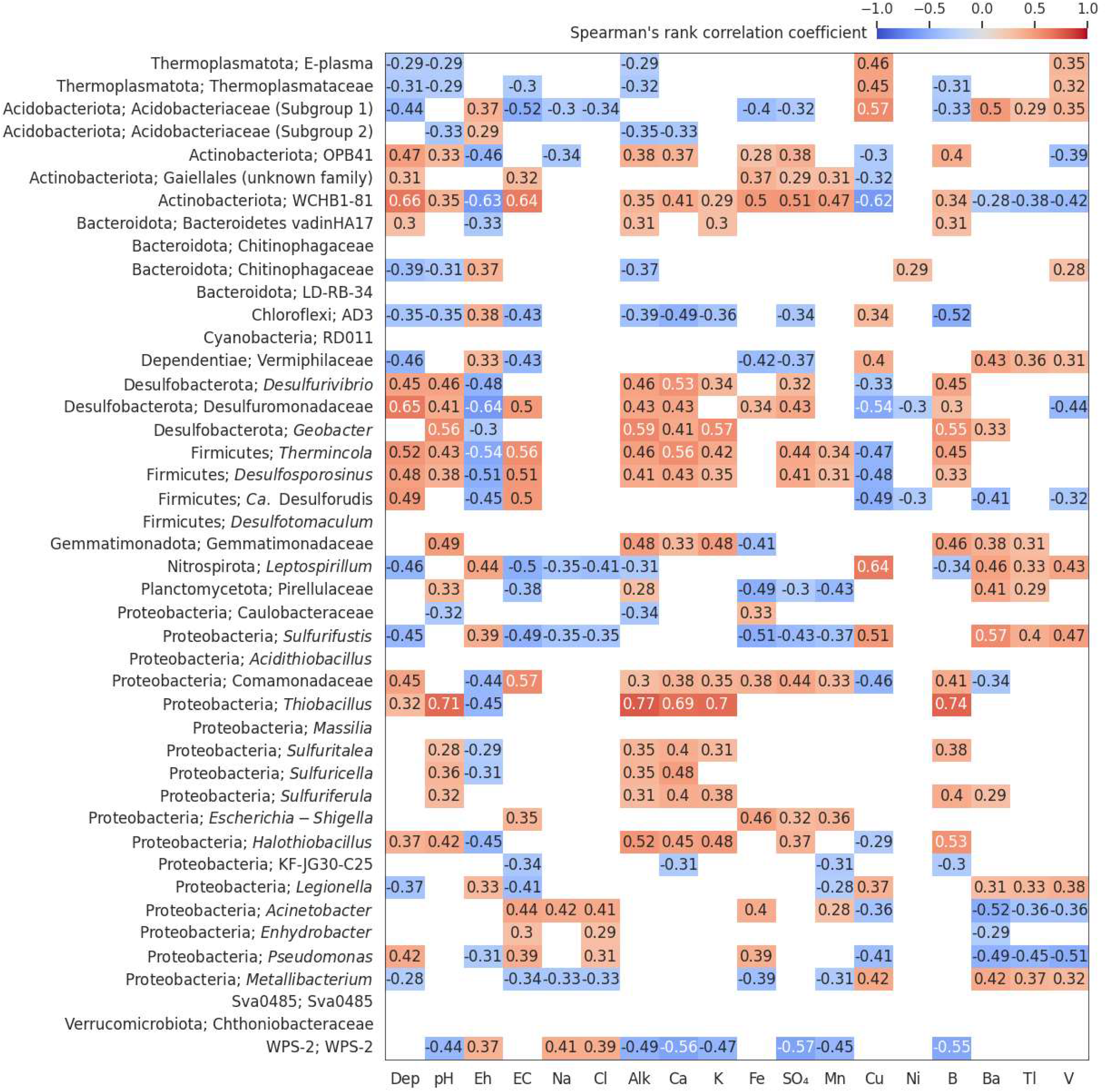
Correlation matrix of genus-level relative abundance and geochemical factors. Spearman’s rank correlation coefficients were calculated between the relative abundance of the abundant genera (Figure 3) and select geochemical factors (Dep = depth, EC = electrical conductivity, Alk = Alkalinity), across all sampling locations. Significant correlations (p<0.05, Bonferroni corrected) are shown. For location-specific correlations, see Supplemental Figure 5.

Diverse consortia associated with iron and sulfur cycling were identified in each core, so their abundance and localization were specifically examined (Figure 3). Iron/sulfur-oxidizing genera common to AMD were typically confined to the upper 2 m of tailings. *Acidithiobacillus* ASVs (the majority of which were classified as *A. ferrooxidans*) were the most abundant predicted iron-oxidizer in the subsurface (0.5 – 1.5 m). The surrounding environment is likely microaerophilic at this depth, which are optimal growth conditions for *A. ferrooxidans* (Namgung & Song, 2015). Pore gas oxygen data was limited to near the tailings surface due to the low gas-filled pore space present in deeper tailings with a high water content and was shown to decrease from atmospheric levels at ground surface to <10% over 1 m in ML25, ML34, and NR18 (Supplementary Figure 4). It is worth noting that *A. ferrooxidans* also exhibits considerable metabolic diversity: not only is this species able to oxidize both ferrous iron and reduced sulfur species in oxic environments, it can also use ferric iron as an electron acceptor in anoxic conditions. Iron reduction is likely the predominant metabolism supporting *A. ferrooxidans* populations under anoxic conditions below depths of 2 metres.

The abundance of *Acidithiobacillus* ASVs in ML34 despite pH > 7 conditions is unusual, but could be associated with the period of exposure to atmospheric oxygen and sulfide oxidation prior to the installation of the DST and organic carbon cover layers. The persistence of *Acidithiobacillus* ASVs could be leveraged for bioremediation approaches, such as in aerobic wetlands that use neutrophilic iron oxidizers to precipitate ferric iron minerals (e.g., ferrihydrite, goethite, magnetite) (Blowes 2003). Existing iron-oxidizers currently applied in aerobic wetlands include *Gallionella* and *Leptothrix* spp., which are not native to AMD (Johnson & Hallberg, 2005). *Acidithiobacillus* species/strains that have adapted to survive in neutral or alkaline environments may be more efficient at immobilizing iron by oxidation while also being tolerant to higher concentrations of copper, nickel, and other toxic metals in tailings.

*Leptospirillum* spp. (mostly *L. ferrooxidans*), exclusive iron-oxidizers, follow a similar distribution but are much less abundant. *L. ferrooxidans* prefers lower pH ranges (0.5-0.7) than *A. ferrooxidans* (1-3) in extreme AMD environments (Schrenk et al., 1998), but was co-localized in moderately acidic environments such as ML25. *A. ferrooxidans* abundance did not significantly correlate with any factor in the combined dataset (Figure 6), but did correlate with a few factors such as pH and Eh within each sampling location (Supplemental Figure 6). *L. ferrooxidans* abundance was moderately-to-strongly correlated with several environmental factors in both the combined dataset and in ML25. Based on these observations, *A. ferrooxidans* could be described as more of a ‘generalist’ species that is able to inhabit many types of mine tailings (see earlier discussion on their metabolic diversity), although they will still localize to a niche within each tailings environment based on preferences in pH, oxygen availability, etc. In contrast, *L. ferrooxidans* is a ‘specialist’ that is adapted to survive at high concentrations of metals (especially copper, R=0.64, although previous studies have shown that their higher affinity for ferrous iron and tolerance to higher concentrations of ferric iron is what drives niche partitioning between *A. ferrooxidans* and *L. ferrooxidans*) (Blowes et al., 2003), but is outcompeted in mine tailings with low metal contamination. It is also important to note that bioleaching with mixed cultures containing both *Acidithiobacillus* spp. and *Leptospirillium* spp. are more effective than pure cultures (Akcil et al., 2007; Falco et al., 2003; Fu et al., 2008; Zhang et al., 2008). Therefore, oxidation and AMD formation within ML25 tailings is expected to be higher due to the combined activity of both organisms (as evidenced by the lower pH and higher dissolved metal content) (Fig 5b), even if their total abundance is similar to single populations of *A. ferrooxidans* at other locations.

Genera predicted to exclusively be sulfur oxidizers included *Sulfurifustis*, *Thiobacillus*, *Halothiobacillus*, *Sulfuritalea*, *Sulfuriferula, Sulfuricella*, and *Desulfovibrio*. The *Sulfurifustis* ASV most abundant in the surface of ML25 and NR18 tailings appears to be a novel species tolerant of acidic environments, as previously isolated *Sulfurifustis* species are neutrophilic (Kojima et al., 2015). The abundance correlation for ASVs in the *Sulfurifustis* genus with geochemical factors resembles that of *Leptospirillum* rather than other sulfur oxidizers, suggesting that members of the *Sulfurifustis* are also specialized for resistance to high metal concentrations (Figure 4). *Thiobacillus* and *Halothiobacillus* ASVs were also not classified to the species level and are predicted to be novel neutrophilic sulfur oxidizers (Whaley-Martin et al. 2019) most abundant in ML34; *Thiobacillus* abundances in particular exhibit a strong positive correlation with pH (R=0.71) and alkalinity (R=0.77). The localization of *Desulfovibrio* (and *Sulfuricella* within NR18) ASVs to the deeper levels of the tailings is in line with their expected use of sulfur oxidation pathways coupled to NO_3_^-^ reduction rather than O_2_ (in the case of *Desulfurivibrio*, it is expected to use the dissimilatory sulfate reduction pathway in the reverse direction), potentially allowing these populations to co-exist with anaerobic sulfate reducers by consuming the sulfides generated by the sulfate reducers (Kojima & Fukui, 2010; Thorup et al., 2017).

The distribution of potential iron reducers was more varied than iron oxidizers, and included *Geobacter*, *Acinetobacter*, *Thermincola, Metallibacterium,* and *Pseudomonas* populations (Pan et al., 2017; Zavarzina et al., 2007; Ziegler et al., 2013). *Geobacter* and *Thermincola* generally localize to greater depths in NR18/NR3, where dissolved iron is high. At pH > 2.5, ferric iron precipitates as secondary oxyhydroxide minerals (Blowes et al. 2003), and reductive dissolution is problematic in these conditions, as it re-mobilizes ferrous iron. The *Pseudomonas* and *Acinetobacter* do not display a consistent localization pattern, but as these genera are highly diverse and ubiquitous in various environments, predictions about iron-reducing capabilities should only be done on the species level, which was not possible as the most abundant ASVs were only classified at the genus level. The *Metallibacterium* are a recently described genus, with one isolate (*M. scheffeleri*) from an acidic biofilm demonstrating iron reduction (Ziegler et al., 2013), although Bartsch et al. (2017) could not identify any iron-reducing genes through genomic, transcriptomic, and proteomic analyses. Unlike the other potential iron reducers, *Metallibacterium* ASVs co-localized with iron oxidizers (*L. ferrooxidans* in ML25, and *A. ferrooxidans* in ML34/NR18); as they are facultative anaerobes and only reduce iron under anoxic conditions (Ziegler et al., 2013), it is unlikely that they are actively reducing iron at these depths. However, as *Metallibacterium* have shown the potential to alkalinize their surroundings through the release of ammonium (Pan et al., 2017; Bartsch et al., 2017), they are a promising candidate for AMD bioremediation and will be a target for investigation in future multi-omic studies.

Sulfate reducers including members of the Firmicutes (*Desulfosporosinus*, *Ca.* Desulforudis, *Desulfotomaculum*) and Desulfobacterota (unclassified Desulfuromonadaceae) were generally dominant at depths >2 m, which is below the depth of oxygen ingress. As expected, the sulfate reducers were generally positively correlated with pH (most are inhibited below pH 5.5) (Hao et al., 1996) and negatively correlated with Eh. They also show moderate negative correlations with dissolved copper concentration, as they are expected to stimulate secondary covellite (CuS) precipitation via the production of sulfide. Relative abundances were not consistent between locations, which could reflect preferences of individual taxa to specific environmental conditions, or legacy impacts from the initial subsurface community present in the ore from which the tailings were generated. The *Desulfotomaculum*, almost exclusively localized to ML25, are a genus mostly associated with deep subsurface environments (Aüllo et al., 2013). *Ca.* Desulforudis (localized to NR18) is also typically found in very deep environments, the most notable being *Ca.* Desulforudis audaxviator, which was the sole organism present 2.8 km below the surface in a gold mine (Chivian et al., 2008). *Desulfosporosinus* are common in natural soil/sediments as well as mine tailings in cold climates (Abicht et al., 2011), and were more evenly distributed in Strathcona Mill tailings compared to Nickel Rim. *Desulfuromonadaceae* abundance followed a similar trend. Finally, the Sva0485 clade (*Ca*. Acidulodesulfobacterales) have recently been described as a dominant member of sulfate-reducing consortia in AMD and ferruginous lakes (Tan et al., 2019). While the characterized members of the Sva0485 lineage possess genes for dissimilatory sulfate reduction, they are facultative anaerobes and also encode genes for sulfur oxidation, iron cycling, and methanogenesis (Tan et al., 2019; Vuillemin et al., 2018). This varied metabolic potential could explain why their depth range was not consistent between tailings locations here, and why their abundance was not significantly correlated to geochemistry (Figure 6), although individual locations showed variable correlations between Sva0458 and salinity (Supplemental Figure 6). Biological sulfate reduction is leveraged in a variety of AMD bioremediation strategies, including anaerobic (or compost) bioreactors, sulfidogenic bioreactors, and permeable reactive barriers (reviewed in Johnson & Hallberg, 2005). All these approaches rely on biogenic sulfide production, typically in the form of H_2_S, which precipitates various metal-sulfide minerals (Blowes et al., 2003; Johnson & Hallberg, 2005).

The archaeon E-plasma (and other unclassified *Thermoplasmataceae* ASVs) abundant at ∼1 m depth at ML25 and NR18 have been identified in various AMD and non-AMD environments (Golyshina, 2011; Baker & Banfield 2003). E-plasma were found to dominate microbial AMD communities within the Parys Mountain mine (UK), characterized by low-to-moderate temperatures and extremely low pH (Korzhenkov et al., 2019), suggesting that they are adapted to colder temperatures (unlike most members of the order *Thermoplasmatales*, which are mesophilic to moderately thermophilic) (Golyshina, 2011). The metabolic potential of E-plasma and other related ‘alphabet-plasma’ lineages belonging to the *Thermoplasmataceae* is currently unknown, as isolation attempts have not yet been successful. However, genomic data suggests that they are heterotrophic scavengers (Korzhenkov et al., 2019).

Many of the most abundant lineages are taxonomically novel and/or previously not identified within mine tailings (Figure 3). The *Actinobacteriota* (most abundant in ML25 but present at all locations) ASVs mostly belong to the order *Gaiellales* (common in deep sea sediments) (Chen et al., 2021), WCHB1-81(microcystin-contaminated lakes) (Dziga et al., 2019), and OPB41 (subsurface environments) (Khomyakova et al., 2022). Other ASVs that were unclassified at the genus/species level include the *Acidobacteriaceae* (associated with mine tailings but highly diverse (Campbell, 2014), as shown by the fact that the two subgroups identified in this study exhibit very different localization patterns), as well as the *Gemmatimonadaceae*, *Pirellulaceae*, *Comamonadaceae*, *Chitinophagaceae*, and *Caulobacteraceae*, which have not previously been associated with mine tailings. The high amount of unresolved microbial diversity from 16S rRNA gene sequencing data implicates that the Strathcona Mill and Nickel Rim mine tailings host currently uncharacterized microorganisms that could participate in the biogeochemical processes governing the release, mobility, and/or attenuation of contaminants associated with AMD. Future multi-omic and culture-based approaches are required to elucidate the full range of microbial activities present at these sites.

### Temporal variation in NR18

Changes in genus-level abundance of ASVs sequenced from NR18 tailings in 2019 compared to 2021 samplings show that the abundance of most genera remained relatively stable over the two-year period (Figure 7). Both *Acidithiobacillu*s and *Pseudomonas* ASVs show an overall trend of decreased abundance at various depths between 0.8 - 2.6 m. We note that prominent increases and decreases in abundance in adjacent core sections is more likely due to imperfect depth mapping between the two years’ cores rather than large changes in community composition at NR18 as a whole. The WPS-2 relative abundance increased at multiple depth ranges between 1-2 m. Many of the NR18 vadose zone pore-water samplers were dry in 2021, and core moisture content is likely to influence WPS-2 abundance, as they are known to be common in dry, bare soil environments (Sheremet et al., 2020). Drier tailings could also explain the decrease in *A. ferrooxidans*, which are not desiccation tolerant (Díaz-Tena et al., 2017). The taxonomic representation of sulfate-reducing bacteria also changes slightly (*Ca.* Desulforudis appear to be superseded by *Desulfosporosinus* between 3-3.5 m), which may be a response to changes in an environmental factor such as pH or alkalinity (moderately positively correlated with *Desulfosporosinus* abundance but not *Ca.* Desulforudis) (Figure 6). Water samples were not collected in 2019, so we were unable to further investigate this hypothesis.

**Figure 7:**
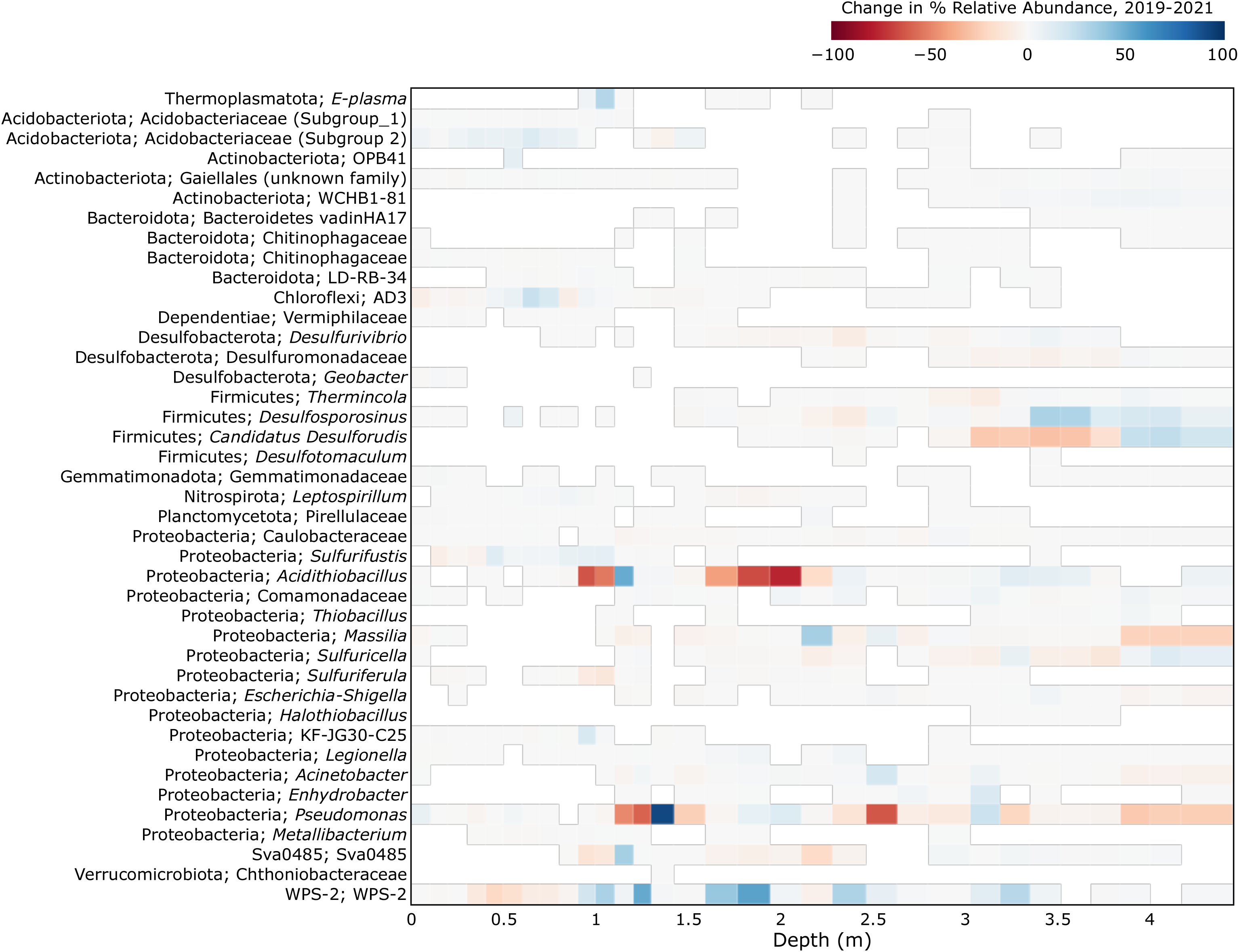
Change in relative abundance of microbial genera across varying depths at the NR18 tailings dump, 2019-2021. The genera listed in Figure 3 were compared to NR18 samples taken in 2019, those present in both datasets were included in this figure. The percent relative abundance of the 2019 samples were subtracted from the NR18 2021 samples. Samples were mapped to each other based on the proximity of the midpoint depths of each core section.

## CONCLUSIONS

Mine tailings present a challenge in waste management practices. AMD environments harbor taxonomically and metabolically diverse microorganisms, which have evolved a wide variety of adaptations to thrive in highly heterogenous contaminated environments. We utilized a 16S rRNA amplicon sequencing approach to characterize microbial communities at narrow (∼10 cm) depth intervals at four locations at two tailings sites, providing a high-resolution profile of community changes across tailings gradients of varying geochemical compositions.

Our research shows that microbial communities are highly diverse between and within each sampling location. Our findings suggest that future sampling efforts that capture a wide range of tailings locations and depths could contribute to the isolation of a broad range of bioremediation-relevant species. A wide variety of microbial guilds are applicable to AMD bioremediation but making improvements in bioreactor/*in-situ* remediation efficiency and cost-effectiveness depends on understanding the appropriate isolates/consortia to leverage given the specific geochemistry of the target mine tailings. We identified that overall community composition is correlated with pH, Eh, alkalinity, salinity, and some metal ions, which surprisingly did not include the major constituents Cu and Ni. Other variables that did not correlate with overall diversity were DOC, F, SO_4_^2-^, NO_3_^-^, Al, As, Co, Li, Mg, Pb, S, Si, Sr, and Zn.

The abundance of most individual lineages was also closely associated with different geochemical factors. However, due to the covarying nature of many environmental variables, it is important to frame predictions about which factors microorganisms are directly responding to, as well as the trajectory of contaminant cycling within tailings, around what is currently known about the physiology and ecology of each individual organism (or its closest known relatives).

Our results showed that amplicon sequencing data can be used to identify specific microenvironments of tailings harboring novel or currently uncultured species, although this approach is limited in its ability to predict the physiological and metabolic properties of these uncharacterized organisms. Many ASVs present in Strathcona Mill and Nickel Rim tailings could not be classified at the species level, including potential iron/sulfur oxidizers and reducers. Functional profiling of tailings known to contain these novel lineages using metagenomics and metaproteomics will be an important future direction to identify genes associated with metal (Fe, Cu, Ni) resistance and transformation. By informing on geochemical variables required to provide niche-specific conditions for growth in the lab our work also provides an insight into improved approaches for establishing enrichment cultures that target these species. The ultimate goal of these combined approaches is to contribute to the development of *in-situ* bioremediation strategies for AMD, or to limit the extent of contaminant release that is accelerated by microbes (Gould et al., 1997).

## ACKNOWLEDGEMENTS

We acknowledge the support of our industry partners at Glencore Sudbury INO for site access, help with field work, and historic site information. We also thank members of Dr. David Blowes and Dr. Carol Ptacek’s research group: Mr. Austin Miller, Ms. Lisa Kester, Mr. YiZhi Yuan, and Mr. Luke Schofield for their help during field sampling. We thank Ms. Emilie Spasov for her help in tailings core processing, and Ms. Rosalind Wang for her help with DNA extractions.

This work was supported by an Ontario Research Fund – Research Excellence grant (ORF-RE09-061). MC was supported by an NSERC PGS-D fellowship. DSG was supported by an NSERC Banting postdoctoral fellowship. LAH and DWB were supported by the Canada Research Chairs.

## METHODS

### Tailings core sampling and processing

Sampling of all locations took place in November 2021. NR18 was previously sampled in November 2019, with the 2019 samples included here for an assessment of temporal variation. In November 2021, the average monthly temperature was -0.5 °C and precipitation was 0.06 mm (Environment and Climate Change Canada, 2022). Sampling in 2019 and 2021 followed the same protocol: Tailings cores of 5.0-7.6 cm (2-3 inches) in diameter were extracted to a depth of approximately 1 meter below the water table (approximately 3-5 m per location) using a similar method as described in (Starr & Ingleton, 1992), with a Pionjar 120 gas-powered hammer drill and aluminum pipe. Cores were capped and stored for 1-2 days on-site before being transported back to the lab, where they were stored at -20 °C prior to further processing.

Cores were sliced into subsamples at 10 cm intervals along the length of each core using a pipe cutter. The centre inch of each subsection was punched out into a sterile plastic bag using a sterile aluminum hollow rod and then stored at -80 °C prior to DNA extraction.

### Geochemical assessments

Pore-water was collected from model 1900 soil water solution samplers (SWSS, from Soil Moisture Inc.) and piezometers installed approximately 1 week prior to sampling. Measurements of pH, Eh, alkalinity, and electrical conductivity were conducted on-site. Additional water samples were filtered (0.45 μm) and stored at 4 °C for laboratory analysis. Inductively coupled plasma-optical emission spectrometry (ICP-OES), inductively coupled plasma-mass spectrometry (ICP-MS) and ion chromatography (IC) were used to determine cation and anion concentrations respectively. Total dissolved organic carbon (DOC) was measured with a non-dispersive infrared sensor (NDIR).

Field measurements of gaseous O_2_/CO_2_ concentrations were taken using the QUANTEK 902P O_2_/CO_2_ analyzer, at approximately 10 cm depth intervals (water saturation of tailings beginning between 1-2 m in depth prevented gas quantification in deeper samples).

### DNA extractions

Approximately 10 g of tailings were measured into a 50 mL falcon tube, washed twice with 35 mL TE buffer (10 mM Tris-HCl, 1 mM disodium EDTA, pH 8.0) to reduce inhibitor concentrations prior to extraction, and transferred to the QIAGEN DNEasy PowerMax Soil kit. DNA was extracted following the manufacturer’s instructions, with the following modifications. DNA was initially eluted with 500 μL of elution buffer (10 mM Tris). Due to low concentrations in 23 samples (10 from NR18 and 13 from NR3), the protocol was altered in subsequent samples. DNA was eluted with 5 mL of elution buffer and subsequently concentrated to a volume of 50-100 μL using ethanol/sodium acetate precipitation as follows: 2.5 volumes of ice-cold ethanol was added to eluted DNA, along with 1/10^th^ volume 3 M sodium acetate. Precipitation reactions were incubated at -20 ℃ overnight, following which precipitated DNA was pelleted at 4000 g and 4 ℃ for 30 minutes, and washed twice with 500 µL of 75% ethanol, spinning for 10 minutes post-wash each time at 21,000 g held at 4 ℃. The pellets were air dried and resuspended in 50-100 uL of elution buffer. Final DNA concentrations were determined using the Thermo Fisher Qubit dsDNA High Sensitivity Assay Kit.

A total of 112 DNA samples were obtained, which were stored at -20 °C prior to sequencing.

### 16S rRNA amplicon sequencing

Samples were PCR-amplified and sequenced by MetagenomBio Life Science Inc. using the Illumina MiSeq machine (v2 reagents). Modified universal primers targeting the V4 region were used for amplification (forward primer 515FB: 5’ GTGYCAGCMGCCGCGGTAA; reverse primer 806RB: 5’ GGACTACNVGGGTWTCTAAT) (Walters et al. 2015).

Sequence data were processed and analyzed using the QIIME2 platform (v.2022.2) (Bolyen et al., 2019). Paired-end reads were filtered, trimmed, merged, and grouped into amplicon sequence variants (ASVs) with DADA2 (Callahan et al., 2016). Taxonomy was assigned based on the SILVA 138 SSU database (Yilmaz et al., 2014). ASV abundance and metadata were processed and visualized in Python (v3.8.8) using the pandas (v1.4.4) (Mckinney, 2010) and matplotlib (v3.6.0) libraries (Hunter, 2007).

### Diversity and statistical analyses

α-diversity was evaluated within QIIME2 (using a rarefied sampling depth of 12,000, which excluded 18 samples) using Faith’s phylogenetic distance, Shannon diversity (H’) index and the observed ASV count. Significant differences between locations were determined using the Kruskal-Wallis test. Plots were generated with Python using the seaborn library (Waskom, 2021).

The QIIME2 feature table and taxonomy data was exported into R (v 4.1.2) (R Core Team, 2021) using the qiime2R package (Bisanz, 2018). Bray-Curtis, weighted and unweighted unifrac distances between samples were calculated to assess β-diversity, using the phyloseq package (McMurdie & Holmes, 2013). Ordination plots were visualized with ggplot2 (Wickham, 2016).

To map each sample to the pore-water chemistry data, the midpoint depth for each core slice was determined (after correcting for the compression factor of the core). Then, the two closest depths (above and below) this value where water chemistry was measured were taken and a weighted average of each geochemical measurement (pH, Eh, dissolved ion concentration, etc.) was calculated based on their respective distance to the midpoint of the core. For dissolved ions, only parameters with ≥50% of measurements above the detection limit of instruments in all cores were included in statistical analyses (with the exception of dissolved organic carbon (DOC) and NO_3_^-^, which were included because they are key metabolic substrates required to support anaerobic respiration and heterotrophy), and values below the detection limit were set to 0.

To determine the geochemical factors affecting β-diversity, a non-metric multidimensional scaling (NMDS) analysis was performed in R using the vegan package (Oksanen, 2007). NMDS ordinations were calculated with the metaMDS function using Bray-Curtis distances. The ‘envfit’ function was used to fit geochemistry factors onto the ordination, using 999 permutations. Variables showing significant correlations (p<0.05, Bonferroni corrected) with community composition were included in the NMDS plot.

Geochemical variables that significantly correlated with community diversity (as well as Cu, Ni, and SO_4_^2-^ concentrations due to their importance as environmental contaminants) were also evaluated for their correlation to the abundance of the most dominant genera (present at a relative frequency of ≥15% in at least one sample). Spearman’s rank correlation matrices were generated using SciPy (v.1.0) (Virtanen et al., 2020) in Python and plotted using the seaborn package. As environmental factors often co-vary, a pairwise correlation matrix between geochemical variables was also calculated.

### Data Availability

All reads associated with amplicon sequencing data presented here have been submitted to the SRA under BioProject PRJNA1017420 and accessions SRR26117691-SRR26117804. All geochemical data used in this study is available in Supplemental Data File 1.

